# Muscle Activation of Upper Body in Different-Angle Suspension Push-Ups: An Analysis of Angle-Specific Muscle Engagement

**DOI:** 10.1101/2025.02.19.638473

**Authors:** Yu-Chin Lin, Kuei-Fu Lin, Chu Chen, Shih-Hsuan Chan, Po-Chun Lee, Chih-Hsiang Hsu, Tsung-Lin Chiang

**Affiliations:** Graduate Institute of Sport Coaching Science, Chinese Culture University, No. 55, Hwa-Kang Rd., Yang-Ming-Shan, Taipei City 11114, Taiwan (R.O.C.); Department of Physical Education, National Tsing Hua University, No. 521, Nanda Rd., East Dist., Hsinchu City 300, Taiwan (R.O.C.); Department of Physical Education and Sport Sciences, National Taiwan Normal University, 2F., No. 1, Aly. 21, Ln. 260, Wuxing St., Xinyi Dist., Taipei City 110, Taiwan (R.O.C.)

**Keywords:** functional training, angle of movement, electromyography

## Abstract

The aim of this study was to understand the effect on upper body muscle activation of bodily angle during push-ups performed with TRX suspension training. Nineteen men (age: 21.1 ± 1.2 years; height: 174.1 ± 4.9 cm; weight: 70.7 ± 7.2 kg) with resistance training experience participated in this study. The participants were required to perform five push-ups on a stable surface and with TRX at five angles (+30°, +15°, 0°, −15°, and −30°, where 0° indicates that the shoulder joints were at the same height as the ankle joints when the arms were extended). The recovery period between the sets at each angle was 3–5 minutes. Stable-surface and TRX push-ups were separated by at least 48 hours. During push-ups, electromyography (EMG) data from the pectoralis major (PM), anterior deltoids (AD), triceps brachii (TRI), upper trapezius (UT), and serratus anterior (SA) were recorded. Muscle activation was indicated by the percentage of maximum voluntary isometric contraction (%MVIC). The %MVIC of each muscle group was then categorized. Repeated-measured two-way analysis of variance was used to determine differences in the muscle group activation between the push-up surface types and five angles. Statistical significance was set at p < .05. The activation of the PM, AD, and TRI during TRX push-ups was categorized as medium to extremely high. Compared with that for stable-surface push-ups, the activation of the PM during TRX push-ups was significantly higher (p < .05). Furthermore, +30°, +15°, and 0° push-ups produced greater PM activation than push-ups at lower angles (p < .05). Both TRX and stable-surface push-ups resulted in greater anterior deltoids, TRI, UT, and SA activation during −30° push-ups. This study indicates the appropriate push-up practices for different muscle groups, as determined by quantifying the muscle activation.

## Introduction

Suspension training system (STS) serves as a functional training apparatus. Its defining feature lies in its utilization of the unstable environment created by the suspension belt, which alters the positioning of the center of mass of the body and the base of support. This alteration consequently enhances muscle activation and motor control capabilities [1, 2]. STS has gained widespread application, encompassing strength training, enhancement of stability, and optimization of motor expression [3, 4]. By utilizing a suspension strap to support the body, STS amplifies the stability challenges during movement, necessitating the collaboration of primary movers and assisting muscle groups with the stabilizing muscle group to uphold stability and precision in movement [5]. Such an unstable environment can intensify the challenges associated with task execution, thereby augmenting strength, stability, and movement expression.

The push-up, a prevalent exercise within the realm of suspension training, serves to effectively engage the pectoralis major (PM), anterior deltoid (AD), triceps brachii (TRI), and the muscle groups responsible for shoulder stabilization (including serratus anterior, SA) [6, 7]. Study evidence indicates that the instability characteristic of the STS elevates the muscular demands for maintaining stable movement, thereby significantly enhancing coordination and presenting an increased challenge [8]. Additionally, engaging in push-ups on a suspension apparatus, as opposed to conventional stable surfaces, has been shown to elicit heightened muscle activation [5], especially within the PM, AD and TRI [9–11]. Moreover, the inherent instability of the suspension apparatus compels the muscular groups to perpetually modify load distribution throughout the exercise, further augmenting coordination and stability of movement.

Changes in body angle significantly influence load adjustment and stability during suspension push-up. The STS modulates intensity via angle and height adjustments of the suspension strap [12]. This modulation is grounded in principles of stability, vector resistance, and pendulum dynamics [1]. Studies reveal that instability demands increased activation of shoulder stabilizers (e.g., serratus anterior, [SA], upper trapezius, [UT]), while major driving muscles (e.g., PM) may exhibit reduced activation due to stability shifts [2]. Additionally, the interaction of suspension strap dynamics and angle variations profoundly affects load distribution and cooperative muscle activation, particularly with elevated foot height. Near-horizontal body angles result in significant activation increases in the triceps and serratus anterior; conversely, raising the foot above shoulder height may alter the activation patterns of the AD and UT [13, 14]. However, the evidence regarding muscle activity at different angles remains insufficient. Nevertheless, the specific activation characteristics of these muscle groups under extreme angle conditions, such as when foot height exceeds shoulder height, remain inadequately explored.

In push-up-related research, comparisons between upper and lower limb elevation have been conducted, with most studies focusing on differences between stable and unstable surfaces [15, 16]. While some investigations have examined the effects of the push-up angle [17] and upper limb elevation [6], the majority of these studies focus on scenarios where the hands and feet are positioned at similar elevations. Limited findings address decline push-ups. In training practice, lower limb elevation is often used as a practical method to increase intensity or introduce variation. However, previous study designs have primarily focused on angles where the shoulder joints are positioned above the lower limbs. Although some studies have explored lower limb elevation during push-ups, the designs of these studies did not involve STS. Hence, the aim of the present study was to determine the muscle activity during push-ups at all possible angles between the torso and ground.

## Methods

### Experimental Approach to the Problem

The present study employed a within-participant repeated-measure design to investigate muscle activity differences during push-ups performed on stable surface and STS (using TRX system). Push-ups were executed at five different angles: the shoulder joints positioned higher (+30° and +15°), lower (−15° and −30°), and parallel (0°) to the ankle joints. muscle activity of the PM, AD, TRI, SA, and UT was recorded throughout the experimental procedure using a surface EMG system (TeleMyo 2400T G2, Noraxon USA). Exercises sequence was counterbalanced. Prior to the experimental trials, MVIC test were performed for each muscle, and the EMG signals were normalized to the respective MVIC values for comparisons across different exercise conditions.

### Participants

Power analysis (GPower, 3.1.9) indicated that a sample size of 13 was required to perform a two-way repeated-measures ANOVA (2*5) with the following input parameters: effect size f = 0.4, alpha = 0.05, power = 0.80, and measurements = 10. Nineteen healthy male university students in physical education were recruited for the study (age: 21.1 ± 1.2 years; height: 174.1 ± 4.9 cm; weight: 70.7 ± 7.2 kg) with at least 6 months resistance training experience and no significant injury to the upper or lower limbs in the previous 6 months. Additionally, the participants were required to have no diagnoses of cardiovascular, musculoskeletal, chronic pulmonary, or arthritic diseases. This study was approved by the National Tsing Hua University Institutional Review Board (No. 10711HS080) and conducted in accordance with the Declaration of Helsinki.

### Procedures

Participants completed two sessions: a familiarization session and an experimental session, both conducted at the same time of day in the morning. The familiarization session took place 48–72 hours before the experimental session to ensure consistency. Participants were required to adhere to specific pre-session guidelines, including refraining from consuming food, beverages, or stimulants (e.g., caffeine) within 2 hours of each session and avoiding vigorous physical activity beyond routine daily activities for at least 24 hours prior to the exercises.

### Familiarization sessions

Each participant was required to participate in two familiarization sessions in which they were shown how to perform TRX push-up at +30°, +15°, 0°, −15°, and −30° (Fig). The angle was defined as that between the straight line from the shoulder joint to the ankle joint and the ground. A horizontal body line represents 0°; +30° and +15° indicate that the shoulder joints were higher than the ankle joints; and −30° and −15° indicate that the shoulder joints were lower than the ankle joints. The purpose of familiarization sessions was to ensure the participants were fully aware of how to perform all movements and capable of performing these movements without any assistance.

The TRX strap length and height of the aerobic steps (which were used to elevate the feet; Fig) were also recorded to determine the appropriate settings for the formal experiment. If a participant was unable to master the push-up movements within the two familiarization sessions, three or more sessions were held until the participant had fully mastered the movements.

**Fig:**
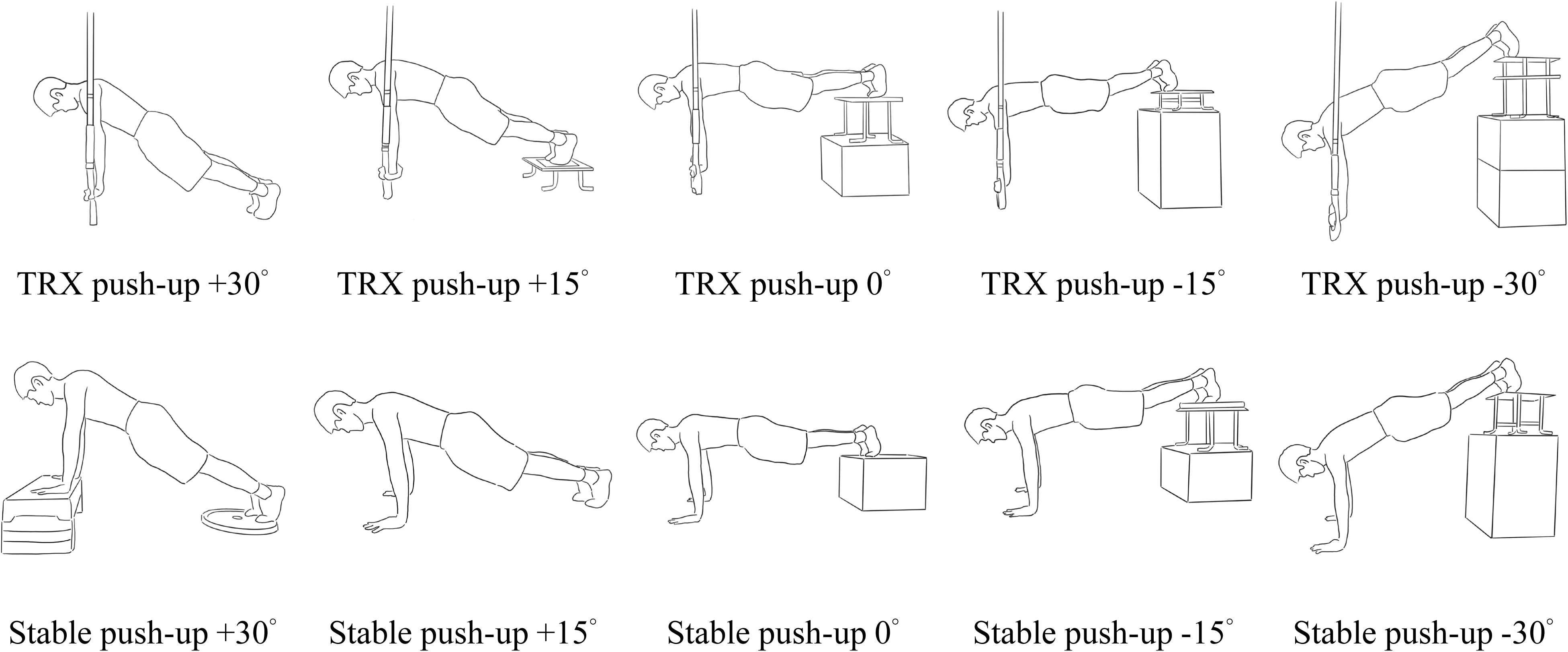
TRX and stable push-up angles

### MVIC determination

For normalization of the EMG data, each participant was required to perform MVIC for each muscle, and EMG responses were collected from the skin surface. Before the electrodes were attached, the skin at the electrode attachment sites was shaved using a razor and cleaned with alcohol to ensure accurate electrode placement and minimize impedance. The MVIC test of the PM, AD, TRI, UT, and SA was three MVIC each lasting 5 seconds. A two-minute interval of rest was allotted between the MVIC test of different muscles. Verbal feedback was provided for each subject for the MVIC procedures. Positions for the MVICs were performed according to standardized procedures, the specific experimental procedures were as follows: 1) PM: the shoulder abducted to 90° and the elbow flexed at 90°, and shoulder horizontal adduction was performed [18]; 2) AD: the shoulder joint was flexed 90°, and shoulder joint flexion was performed [18]; 3) TRI: sitting position. the shoulder and elbow joints were flexed 90°, and the forearm pronated, and elbow extension was performed [18]; 4) UT: the shoulder abducted to 90° with the neck side-bent, rotated to the opposite side, and extended [19]; 5) SA: supine position. the shoulder flexed to 125°, and the scapula was upwardly rotated with the shoulder flexed [19].

### Exercise sessions and applications

In the formal experiment, the TRX strap was connected to an anchor point and was perpendicular to the ground. Each participant was required to perform push-ups at five angles (+30°, +15°, 0°, −15°, and −30°). For each angle, five repetitions were performed. The participant had a 3–5-minute recovery interval between sets for each angle. The amplitude of the EMG root mean square (RMS) for the PM, AD, TRI, UT, and SA was recorded during the entire procedure. In all trials, the greatest RMS value recorded during push-ups was averaged and normalized to the MVIC (%MVIC). A previous study’s categorization of muscle activation into four levels was adopted in this study to compare muscle activity levels under different conditions: low (L) = 0–20% MVIC; mid-level (M) = 21–40% MVIC; high (H) = 41–60% MVIC; and very high (VH) = ≥61% MVIC [20].

### Instrumentation and data processing

The muscle activity of the PM, AD, TRI, SA, and UT was recorded using an EMG system (TeleMyo 2400T G2, Noraxon USA) with a sampling rate of 1,000 Hz. Following the recommendations of the Surface EMG for the Non-Invasive Assessment of Muscles (SENIAM) [21], pairs of bipolar surface electrodes were aligned along the muscle midline and oriented parallel to the muscle fibers. The center-to-center distance between electrodes was set at 2.5 cm. Prior to electrode placement, the skin was prepared by shaving and cleaning with alcohol wipes to reduce impedance. A reference electrode was affixed to the acromioclavicular joint, and interelectrode resistance was verified to remain below 10 kΩ for all participants. The collected EMG signals were digitized using the TeleMyo 2400T G2 system and subsequently processed with Myoresearch XP 1.07 software (Noraxon USA Inc.). Signal processing included rectification, band-pass filtering, and integration. EMG amplitude was determined as follows: 1) during MVIC tests, the root mean square (RMS) of the signal was calculated over a 500-millisecond window centered on the peak force; 2) during push-up exercises, the RMS was calculated for each repetition within a 500-millisecond window centered on the highest signal value.

### Data analysis and statistics

Statistical analyses were performed using SPSS version 27.0 for Windows. All data are presented as a mean (M) and standard deviation (SD). The Shapiro–Wilk test was used to assess the normality of the variables. A two-way repeated-measures analysis of variance (ANOVA) was used to compare the differences in muscle activity during push-ups performed on two surfaces (stable surface vs. TRX) and fives angles (+30°, +15°, 0°, -15°, and -30°). If an interaction approached significance, Tukey’s post hoc test and paired t-test was used. The significance level was set at 0.05.

## Results

The muscle activation during push-ups on a stable surface and with TRX at five angles is shown in Table 1. For PM muscle activation, the two-way repeated measures ANOVA showed no significant surface and angles interaction effect (F=1.069, *p* =.364, partial *η²* =.056). However, significant main effects were found for surface (F=20.782, *p* <.001, partial *η²* = .536) and angles (F=19.109, *p* <.001, partial *η²* = .515). Post hoc analysis revealed that PM activation was significantly greater during TRX push-ups than stable-surface push-ups. Regarding angle differences, PM activation at +30°, +15°, 0°, and −15° was significantly greater than at −30°, and at 0° was significantly greater than at −15°.

**Table 1.**
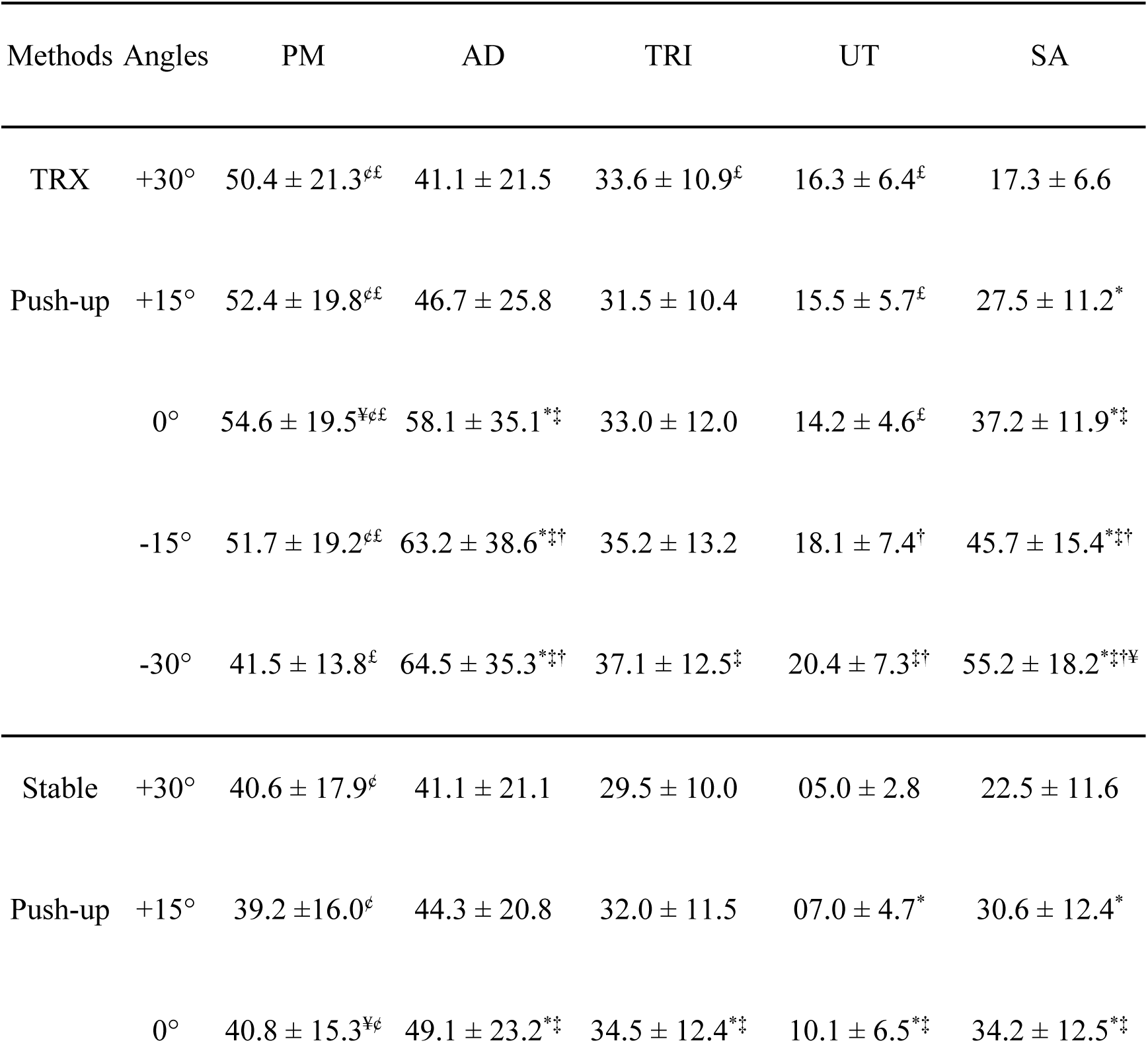
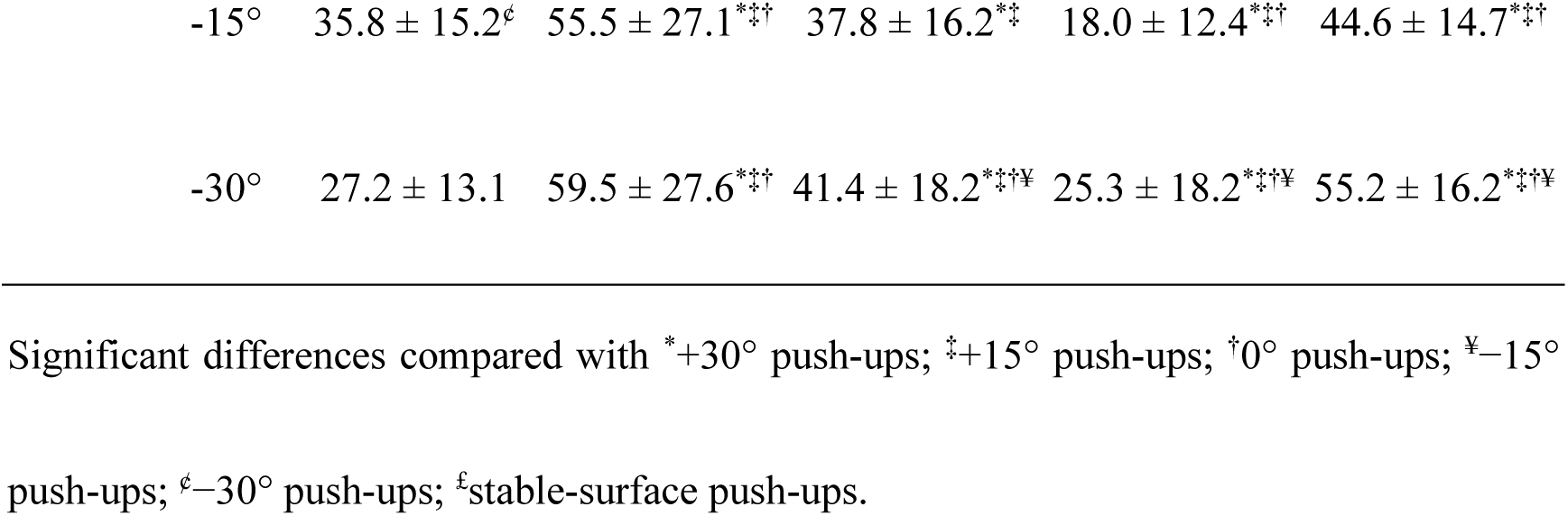
Differences in muscle activity during push-ups with TRX and on a stable surface.

No significant interaction was observed between surface and angles for AD muscle activation (F = 1.581, *p* = .218, partial *η²* =.081). Similarly, no significant main effect was found for surface (F = 1.637, *p* = .217, partial *η²* =.083), but a significant main effect was observed for angles (F = 17.831, *p* < .001, partial *η²* =.498). Post hoc analysis indicated that AD at 0°, −15°, and −30° was significantly greater than at +30° and +15°, while at −15° and −30° was significantly greater than at 0°.

For TRI muscle activation, a significant interaction between surface and angle was observed (F = 4.894, *p* = .016, partial *η²* =.214). Post hoc comparisons revealed that during TRX push-ups, TRI was significantly greater at −30° than at +15° (F = 5.405, *p* = .007, partial *η²* =.231). On a stable surface, TRI at 0°, −15°, and −30° was significantly greater than at +30° and +15°, and at −30° was significantly greater than at 0° and −15° (F = 14.727, *p* < .001, partial *η²* =.450). Moreover, at +30°, TRI was significantly greater during TRX push-ups than during stable-surface push-ups (*p* = .023, *d* = -.571, 95% CI [-1.051, -.078]).

A significant interaction between surface and angle was also observed for UT muscle activation (F = 10.432, *p* = .001, partial *η²* =.367). Post hoc analysis revealed that during TRX push-ups, UT at −15° and −30° was significantly greater than at 0°, and at −30° was significantly greater than at +15° (F = 6.043, *p* < .001, partial *η²* =.251). On a stable surface, UT activation differed significantly across angles (F = 23.716, *p* < .001, partial *η²* = .569). Post hoc comparisons indicated significant differences between all angles (p < .05), with activation highest at −30°, followed by −15°, 0°, +15°, and lowest at +30°. Moreover, UT activation was significantly greater during TRX push-ups compared to stable-surface push-ups at +30° (*p* < .001, *d* = -1.728, 95% CI [- 2.437, -1.001]), +15° (*p* < .001, *d* = -1.692, 95% CI [-2.386, -0.980]), and 0° (*p* = .031, *d* = -1.370, 95% CI [-1.993, -0.728]).

Finally, a significant interaction between surface and angle was observed for SA muscle activation (F = 2.913, *p* = .027, partial *η²* =.139). Within-condition analyses revealed that SA activation significantly differed across angles in both TRX push-ups (F = 109.850, *p* < .001, partial *η²* = .859) and stable-surface push-ups (F = 79.535, *p* < .001, partial *η²* = .815). Post hoc comparisons showed significant differences between all angles, with SA activation highest at −30°, followed by −15°, 0°, +15°, and lowest at +30° in both conditions. No significant differences between the different surfaces.

## Discussion

Previous studies have quantified intensity and load using TRX straps (with a force meter) and the ground reaction force; however, clearly discriminating the loads bore by the upper and lower limbs, respectively, was difficult [22, 23]. This study analyzed muscle activity using EMG and employed the %MVIC as a quantitative indicator.

According to Table 1, as the push-up angle changed, muscle activation varied. When the position of the shoulder joints was higher than (or the same height as) the ankle joints (+30°, +15°, and 0°) during push-ups, the PM were greatly activated. When the shoulder joints were lower than the ankle joints (−15° and −30°), the muscle activity was lower. Conversely, the AD, TRI, UT, and SA showed higher muscle activity when the shoulder joints were positioned lower than the ankle joints, with the greatest activation occurring at −30°.

As the TRX push-up angle decreases, significant changes in upper limb muscle activation patterns are observed. Our findings indicate that activation of the AD, TRI, UT, and SA significantly increases (Table 1). Studies have demonstrated that lower TRX push-up angles shift a greater proportion of body mass load onto the upper limbs, thereby increasing the recruitment of these muscle groups [17, 22, 23]. In contrast, the activation pattern of the PM exhibits a different trend. At +30°, +15°, and 0°, its activation remains relatively high; however, as the angle decreases to –15° and –30°, activation declines. This change may be attributed to increased shoulder flexion with decreasing TRX push-up angles, which alters the primary muscle recruitment pattern and reduces the contribution of the PM. These findings indicate that variations in suspension push-up angles affect both load distribution and the relative contributions of different muscle groups.

In previous studies, the feet have been placed beneath the anchor point of suspension or the relative position of the feet to the anchor point has been altered to approach the target angle [6, 16, 22, 23]. However, this study standardized the TRX anchor point and positioned the arms and straps directly beneath it. (Fig). This design reduced potential confounding factors, such as shifting body weight or variations in relative positioning, ensuring that angle variation was the primary factor influencing muscle activation. This design also allowed for a greater range of angle variations in TRX exercises, making it more advantageous for training applications.

Although previous studies have reported that TRX instability enhances muscle activation [11, 24, 25]. However, our results indicate that at push-up angles of -15° and -30°, only PM activation was significantly greater in TRX push-ups compared to the stable surface, while other muscle groups exhibited no significant surface-related differences. This suggests that the increased PM activation at lower angles may result from additional force demands required for movement stabilization in the TRX condition. Given that the arms and straps were positioned directly beneath the anchor point in this study, the potential instability effects of the TRX system may have been mitigated, allowing angle variation to play a more dominant role in modulating muscle activation. To further examine how muscle activation varies with TRX push-up angles, we systematically analyzed the activation levels of upper-body muscles across different conditions.

The present study demonstrates that push-up angle plays a more dominant role in modulating upper-body muscle activation than TRX instability, with distinct recruitment patterns across different angles (Table 2). At higher angles (+30°, +15°, 0°), PM activation remained high level, while activation of AD, TRI, SA, and UT was relatively stable. In contrast, at lower angles (-15°, -30°), AD and SA activation increased significantly, while TRI activation showed an unexpected pattern, remaining mid-level across all TRX push-ups but increasing to high level only at -30° on the stable surface. These results suggest that the combined effects of body positioning, force distribution, and joint stabilization demands contribute to these activation differences. Analysis of %MVIC variations (Table 3) further supports these findings, revealing that when the push-up angle was less than 0°, activation of AD, TRI, UT, and SA gradually increased, regardless of whether push-ups were performed with TRX or on the stable surface. Notably, PM activation at -30° was 24% and 33% lower compared to 0° in TRX and stable surface push-ups, respectively, confirming that increased shoulder flexion shifts mechanical demands away from PM. Conversely, UT and SA activation were 1.4 and 1.6 times higher at -30° than at 0°, highlighting their critical role in shoulder stabilization at lower angles.

**Table 2.**
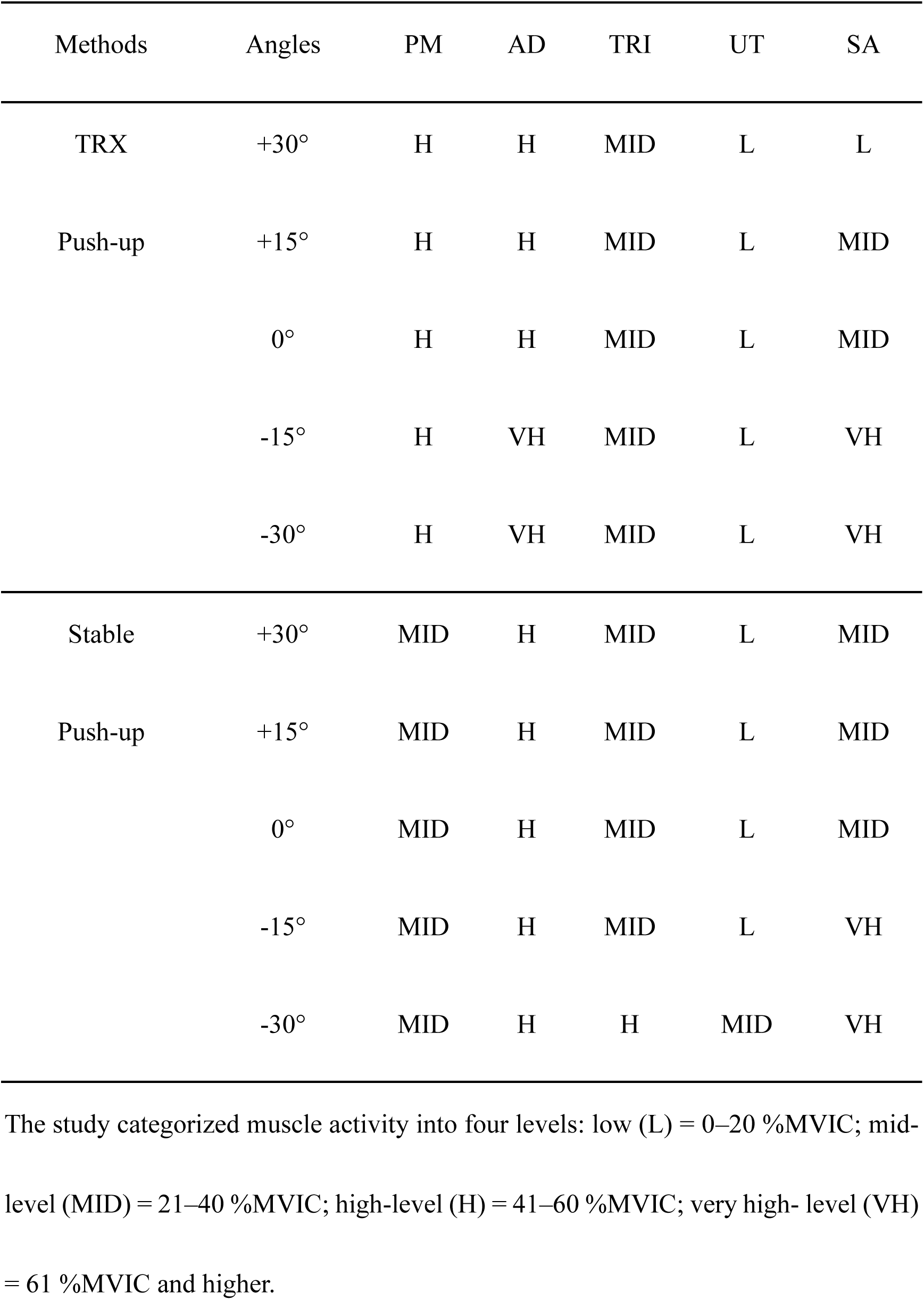
Level of muscle activity during push-ups with TRX and on a stable surface.

| Methods | Angles | PM | AD | TRI | UT | SA |
| --- | --- | --- | --- | --- | --- | --- |
| TRX | +30° | H | H | MID | L | L |
| Push-up | +15° | H | H | MID | L | MID |
|  | 0° | H | H | MID | L | MID |
|  | -15° | H | VH | MID | L | VH |
|  | -30° | H | VH | MID | L | VH |
| Stable | +30° | MID | H | MID | L | MID |
| Push-up | +15° | MID | H | MID | L | MID |
|  | 0° | MID | H | MID | L | MID |
|  | -15° | MID | H | MID | L | VH |
|  | -30° | MID | H | H | MID | VH |

The study categorized muscle activity into four levels: low (L) = 0–20 %MVIC; mid-level (MID) = 21–40 %MVIC; high-level (H) = 41–60 %MVIC; very high- level (VH) = 61 %MVIC and higher.

**Table 3.** Percentage changes in muscle activity in TRX and stable-surface push-ups between the different angles.

|  | PM |  | AD |  | TRI |  | UT |  | SA |  |
| --- | --- | --- | --- | --- | --- | --- | --- | --- | --- | --- |
| Angles | TRX | Stable | TRX | Stable | TRX | Stable | TRX | Stable | TRX | Stable |
| 0°—+30° | -8 | -1 | -29 | -16 | 2 | -15 | 15 | -50 | -53 | -34 |
| 0°—+15° | -4 | -4 | -20 | -10 | -5 | -7 | 9 | -31 | -26 | -10 |
| 0°—-15° | -5 | -12 | 9 | 13 | 6 | 9 | 27 | 78 | 23 | 30 |
| 0°—-30° | -24 | -33 | 11 | 21 | 12 | 20 | 44 | 151 | 48 | 61 |

These variations in mechanical demands appear to drive the observed differences in muscle activation. At lower angles (-15°, -30°), greater shoulder flexion likely increases reliance on AD and SA for stabilization, reducing the necessity for TRI to contribute significantly to movement execution, particularly in TRX push-ups where instability further increases postural control demands. The fixed TRX anchor point may have further influenced this redistribution, limiting TRI engagement by promoting compensatory activation of other stabilizers. In contrast, on the stable surface at -30°, the absence of instability may have necessitated greater elbow extension force, leading to increased TRI activation. These findings suggest that TRX instability does not inherently enhance activation across all muscle groups but instead redistributes neuromuscular effort depending on movement constraints.

While TRX push-ups increased PM activation across all angles, they did not significantly alter TRI, SA, or UT activation compared to stable push-ups. Higher-angle TRX push-ups (+30°, +15°, 0°) primarily target PM, whereas lower angles (-15°, -30°) shift activation toward AD and SA. Given that no single push-up variation maximizes activation across all muscle groups, training programs should adjust angles to target specific muscles rather than relying solely on TRX instability. Future research should examine how joint loading strategies, muscle coordination, and external support conditions interact to optimize neuromuscular control in suspension training.

## Conclusions

Push-up angles and surface conditions should be carefully selected based on target muscle groups and training goals, as muscle activation patterns vary with angle adjustments and stability demands. If the main objective is to strengthen the PM, we suggest that beginners start with push-ups on a stable surface, then increasing the challenge with TRX with the shoulder joints higher than or at equal height to the ankle joints, where the greatest PM activation occurs near 0°. Conversely, if the goal is to strengthen the AD, TRI, UT, or SA, we suggest that beginners start with +30° push-ups on a stable surface and then gradually decrease the angle to increase the intensity, with the highest intensity reached at −30°. Understanding that high activation of the PM results in low activation of other muscle groups is crucial. The lower the position in which push-ups are performed, the less the PM are involved. In terms of overall muscle engagement, TRX push-ups—regardless of angle—may enhance neuromuscular demand more effectively than stable surface push-ups, making them a valuable tool for progressive strength development.

## Notes

### Competing Interest Statement

The authors have declared no competing interest.

## References

1. Bettendorf B. TRX suspension training bodyweight exercises: scientific foundations and practical applications. San Francisco: Fitness Anywhere Inc. 2010.

2. Behm D, Colado JC. The effectiveness of resistance training using unstable surfaces and devices for rehabilitation. International Journal of Sports Physical Therapy. 2012;7(2):226–41. PubMed PMID: 22530196; PubMed Central PMCID: PMCPMC3325639.

3. Soligon SD, da Silva DG, Bergamasco JGA, Angleri V, Júnior RAM, Dias NF, et al. Suspension training vs. traditional resistance training: effects on muscle mass, strength and functional performance in older adults. European Journal of Applied Physiology. 2020;120(10):2223–32. Epub 20200722. doi: 10.1007/s00421-020-04446-x. PubMed PMID: 32700098.

4. Ma X, Sun W, Lu A, Ma P, Jiang C. The improvement of suspension training for trunk muscle power in Sanda athletes. Journal of Exercise Science and Fitness. 2017;15(2):81–8. Epub 20171008. doi: 10.1016/j.jesf.2017.09.002. PubMed PMID: 29541137; PubMed Central PMCID: PMCPMC5812878.

5. Harris S, Ruffin E, Brewer W, Ortiz A. Muscle activation patterns during suspension training exercises. International Journal of Sports Physical Therapy. 2017;12(1):42–52. PubMed PMID: 28217415; PubMed Central PMCID: PMCPMC5294946.

6. Borreani S, Calatayud J, Colado JC, Tella V, Moya-Nájera D, Martin F, et al. Shoulder muscle activation during stable and suspended push-ups at different heights in healthy subjects. Physical Therapy in Sport. 2015;16(3):248–54. Epub 20141218. doi: 10.1016/j.ptsp.2014.12.004. PubMed PMID: 25882770.

7. Kowalski KL, Connelly DM, Jakobi JM, Sadi J. Shoulder electromyography activity during push-up variations: a scoping review. Shoulder and Elbow. 2022;14(3):326–40. Epub 20210606. doi: 10.1177/17585732211019373. PubMed PMID: 35599715; PubMed Central PMCID: PMCPMC9121296.

8. Aguilera-Castells J, Buscà B, Fort-Vanmeerhaeghe A, Montalvo AM, Peña J. Muscle activation in suspension training: a systematic review. Sports Biomech. 2020;19(1):55–75. Epub 20180614. doi: 10.1080/14763141.2018.1472293. PubMed PMID: 29902124.

9. Beach TA, Howarth SJ, Callaghan JP. Muscular contribution to low-back loading and stiffness during standard and suspended push-ups. Human Movement Science. 2008;27(3):457–72. Epub 20080324. doi: 10.1016/j.humov.2007.12.002. PubMed PMID: 18362038.

10. Calatayud J, Borreani S, Colado JC, Martin F, Rogers ME. Muscle activity levels in upper-body push exercises with different loads and stability conditions. The Physician and Sportsmedicine. 2014;42(4):106–19. doi: 10.3810/psm.2014.11.2097. PubMed PMID: 25419894.

11. Snarr RL, Esco MR. Electromyographic comparison of traditional and suspension push-ups. Journal of Human Kinetics. 2013;39:75–83. Epub 20131231. doi: 10.2478/hukin-2013-0070. PubMed PMID: 24511343; PubMed Central PMCID: PMCPMC3916913.

12. Giancotti GF, Fusco A, Varalda C, Capelli G, Cortis C. Evaluation of training load during suspension exercise. The Journal of Strength and Conditioning Research. 2021;35(8):2151–7. doi: 10.1519/jsc.0000000000003100. PubMed PMID: 30893278.

13. Lear LJ, Gross MT. An electromyographical analysis of the scapular stabilizing synergists during a push-up progression. The Journal of Orthopaedic and Sports Physical Therapy. 1998;28(3):146–57. doi: 10.2519/jospt.1998.28.3.146. PubMed PMID: 9742471.

14. Patselas T, Karanasios S, Sakellari V, Fysekis I, Patselas MI, Gioftsos G. EMG activity of the serratus anterior and trapezius muscles during elevation and push up exercises. Journal of Bodywork and Movement Therapies. 2021;27:247–55. Epub 20210304. doi: 10.1016/j.jbmt.2021.02.002. PubMed PMID: 34391241.

15. Lehman GJ, MacMillan B, MacIntyre I, Chivers M, Fluter M. Shoulder muscle EMG activity during push up variations on and off a Swiss ball. Dynamic Medicine. 2006;5:7. Epub 20060609. doi: 10.1186/1476-5918-5-7. PubMed PMID: 16762080; PubMed Central PMCID: PMCPMC1508143.

16. McGill SM, Cannon J, Andersen JT. Analysis of pushing exercises: muscle activity and spine load while contrasting techniques on stable surfaces with a labile suspension strap training system. The Journal of Strength and Conditioning Research. 2014;28(1):105–16. doi: 10.1519/JSC.0b013e3182a99459. PubMed PMID: 24088865.

17. Melrose D, Dawes J. Resistance characteristics of the TRX^TM^ suspension training system at different angles and distances from the hanging point. Journal of Athletic Enhancement. 2015;4(1):2–5.

18. Freeman S, Karpowicz A, Gray J, McGill S. Quantifying muscle patterns and spine load during various forms of the push-up. Medicine and science in Sports and Exercise. 2006;38(3):570–7. doi: 10.1249/01.mss.0000189317.08635.1b. PubMed PMID: 16540847.

19. Ekstrom RA, Soderberg GL, Donatelli RA. Normalization procedures using maximum voluntary isometric contractions for the serratus anterior and trapezius muscles during surface EMG analysis. Journal of Electromyography and Kinesiology. 2005;15(4):418–28. Epub 20041225. doi: 10.1016/j.jelekin.2004.09.006. PubMed PMID: 15811612.

20. Cugliari G, Boccia G. Core muscle activation in suspension training exercises. Journal of Human Kinetics. 2017;56:61–71. Epub 20170315. doi: 10.1515/hukin-2017-0023. PubMed PMID: 28469744; PubMed Central PMCID: PMCPMC5384053.

21. Hermens HJ, Freriks B, Disselhorst-Klug C, Rau G. Development of recommendations for SEMG sensors and sensor placement procedures. Journal of Electromyography and Kinesiology. 2000;10(5):361–74. doi: 10.1016/s1050-6411(00)00027-4. PubMed PMID: 11018445.

22. Giancotti GF, Fusco A, Varalda C, Capranica L, Cortis C. Biomechanical analysis of suspension training push-up. The Journal of Strength and Conditioning Research. 2018;32(3):602–9. doi: 10.1519/jsc.0000000000002035. PubMed PMID: 29466266.

23. Gulmez I. Effects of angle variations in suspension push-up exercise. The Journal of Strength and Conditioning Research. 2017;31(4):1017–23. doi: 10.1519/jsc.0000000000001401. PubMed PMID: 26950344.

24. Borreani S, Calatayud J, Colado JC, Moya-Nájera D, Triplett NT, Martin F. Muscle activation during push-ups performed under stable and unstable conditions. Journal of Exercise Science and Fitness. 2015;13(2):94–8. Epub 20150902. doi: 10.1016/j.jesf.2015.07.002. PubMed PMID: 29541105; PubMed Central PMCID: PMCPMC5812863.

25. Batista GA, Beltrán SP, Passos M, Calixtre LB, Santos LRH, de Araújo RC. Comparison of the electromyography activity during exercises with stable and unstable surfaces: a systematic review and meta-analysis. Sports. 2024;12(4):111. Epub 20240418. doi: 10.3390/sports12040111. PubMed PMID: 38668579; PubMed Central PMCID: PMCPMC11055131.

